# Network-level containment of single-species bioengineering

**DOI:** 10.1101/2021.07.05.451200

**Authors:** Victor Maull, Ricard Solé

## Abstract

Ecological systems are facing major diversity losses in this century due to Anthropogenic effects. Habitat loss, overexploitation of resources, invasion and pollution are rapidly jeopardising the survival of whole communities, as revealed by pronounced population losses. Moreover, the potential of future tipping points further complicate their survival and change our perspective of risk. It has been recently suggested that a potential approach to flatten the curve of species extinction and prevent catastrophic shifts would involve the engineering of one selected species within one of these communities, aiming at helping the maintenance of key conditions compatible with high diversity. Such possibility has started to become part of potential intervention scenarios to preserve coral reefs, kelp forests or soil microbiomes in drylands. Despite its potential, very little is known about the actual dynamic responses of complex ecological networks to the introduction of a synthetic strains derived from a resident species. In this paper we address this problem by modelling the response of a competitive community to the addition of a synthetic strain derived from a member of a stable ecosystem. We show that the community interaction matrix largely limits the spread of the engineered strain, thus suggesting that species diversity acts as an ecological firewall. Implications for future restoration and terraformation strategies are discussed.

## I. INTRODUCTION

Our biosphere is facing multiple challenges associated to global warming, demographic growth and unsustainable economic practices [1, 2]. A rapid loss of biodiversity, both at the population and species levels is a known, an accelerated outcome of the Anthropocene. This is illustrated by recent studies that reveal dramatic decays in insect populations as a consequence of combined effects of habitat loss, pesticides and climate change [3, 4]. Their decay, as well as the loss of species in many other groups, will have deep consequences for the stability of ecosystems. The problem is exacerbated by the possibility that such declines occur following rapid community shifts [5].

Our place in this threatened biosphere is far from disconnected from our fate. There is an urgent need for a shift of perspective that requires changes in ecosystem management on all scales as well as new ways of dealing with Anthropogenic impacts. Social end economic factors are not out of this equation and the potential for ecosystem collapse must be addressed from different approximations that properly weight conservation and human wellbeing.

An example of such situation is provided by drylands. They represent 45% of emerged lands, host around 40% human population and will experience diverse transitions associated with the change in several key functional and structural attributes [6]. These changes will cause shifts and abrupt decays in plant productivity, soil fertility and plant diversity and cover. Can they be counterbalanced by restoration strategies, including the use of genetic engineering? Can positive feedback loops be promoted in endangered communities to preserve them from collapse? [7, 8]

The potential use of genetically modified species has been largely ignored (and sometimes harshly criticised) by most researchers involved in ecosystem restoration. One reason for this (when comparing with other approaches) is the fear that deployment of modified organisms can have unintended consequences. This was the case of the well-known single-gene modification of a microorganism (*Pseudomonas syringae*) that was known to cause the formation of ice crystals on the surface of some plant crops. As a consequence, frost develops on plant buds and crops are lost. The engineering approach was simple: take the extant bacterium and remove one gene responsible for the “ice-plus” protein causing the formation of ice, and spray the plants with it. Despite its success, and the fact that the engineered strain was in fact one possible natural mutant of the wild type strain, almost immediate lawsuits blocked further developments in the area [10]. A similar situation affects the potential efforts for using bioremediation strategies. Bioremediation studies have been developed since the mid 1980s, but largely limited to in *vitro* conditions due to safety issues and variable success [11, 12].

The banning of GMOs created a knowledge gap and many untested assumptions that still prevents to properly address the question of how modified strains will or not affect ecosystems networks [13]. Too often, the impact of (potential) synthetic modified bacteria is compared with the role played by exotic invaders [14–16] that are known to have potential damaging consequences. But the comparison is flawed in many ways. On the one hand, modified organisms are very likely to fail surviving in a given environment outside the laboratory conditions where they have been grown. It has been known since the 1980s that a modified microorganism that has the same genome that a given, resident species, is likely to fail performing their functions or survive. This is a consequence of stresses (biotic responses, niche context, weather conditions) that differ from those used in the lab. However, in recent years some voices are rising in support to synthetic environmental engineering possibilities. One of the reasons is that, along with the emergency situation affecting some endangered ecosystems, the new field of synthetic biology has shown the potential for targeting complex problems in health and disease [17, 18]. More recently, it has reactivated the debate on the use of this genetic engineering to tackle current and urgent ecological problems [19–23]. Two relevant examples include the engineering of synthetic microbiomes for coral reefs [24–26] or Kelp Forest communities [27]. As pointed out by several authors, the interventions are not necessarily targeting the recovery of a previous state [19, 28] but should protect or even enhance species diversity.

**FIG. 1:**
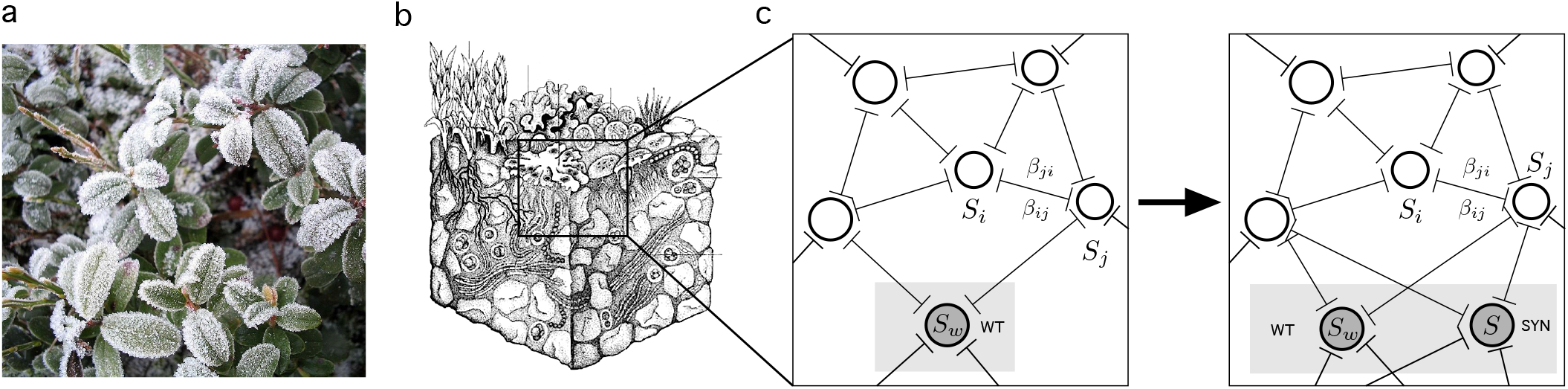
Single-species engineering has been used to treat specific problems in crops, such as ice formation in leaves (a) that can kill plants. Using a modified bacterium lacking a specific gene, ice formation is inhibited. Single-species modifications of this kind could be used to modify the soil microbiome in dryland soil crusts (Figure (b) adapted from [9]). In (c-d) we struck the single-species engineering analyses here. A given, wild type species as *S_w_*, with replication rate *ρ_w_* in the original community is used to create a synthetic strain (d) *S_s_* with replication rate *ρ_s_* > *ρ_w_*.

In this paper we aim at providing some rationale for what could be expected to happen under a well defined set of conditions involving a deployed synthetic microorganism in a given multi species community. Following a recent proposal [19], the idea is to consider a given native species that is used to create a new strain including a slight targeted modification, to be reintroduced in the same community. Although some small-species case scenarios have been previously studied [29, 30]. We take a population dynamics approach where a set of standard species competition models are used to test the behavior of this particular bioengineering approach. The model allows to study the resulting stability and how such introduction affects the community. As will be shown below (with both analytic and numeric arguments) the introduced species gets integrated in the community and no impact in terms of stability is found, thus suggesting that (under these conditions) a diverse ecosystem can intrinsically define an effective population firewall.

## II. METHODS

### A. Multispecies competition model

In this paper we make use of *S*-dimensional Lotka-Volterra (LV) models of competition as a minimal description of a complex community to described both the original and the engineered scenarios [31, 32]. Specifically, the equations read:

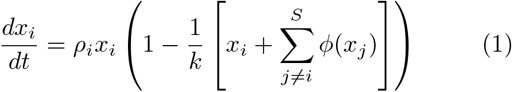

where *x_i_*(*i* = 1, 2, 3…*S*) indicate the population abundance of each species and *ϕ*(*x_j_*) indicates the functional response, defined as *ϕ*(*x_j_*) = *β_ij_x_j_* for the simplest, linear case. Here replication rates *ρ_i_* and the community matrix *β_ij_* are constants with different biological meanings which introduce feedback loops and interactions among species. All species have intrinsic replication rates (*ρ* ∈ (0, 0.3) values) and the community matrix *β* = (*β_ik_*) that will weight the strength of the pairwise interactions [33]. Here, the interspecific, off-diagonal, terms of interaction matrix come up from a uniform rectangular distribution from a uniform distribution *U*[0,1] [34] and *U*[0, 2] in order tot test stronger interactions, see supplementary material. The intraspecific, diagonal terms, are set to *β_ii_* = 1. Furthermore, in order to guarantee stationary attractors, we use *β_ij_* = *β_ji_* [35]. This scenario might reflect the structure of species sharing a common resource, where competition could be more or less symmetrical [36, 37]. Finally the carrying capacity is expressed in *k_i_* = 1 terms.

Along with the linear case, we will also consider Holling type II and type III functional responses [38, 39] that incorporate two main forms of saturation functions. Specifically, we will have

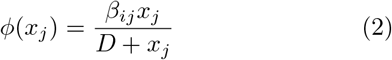

for Holling type II: and

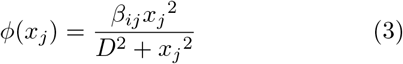

for Holling type III, respectively. Here *D* stands as the half-saturation density of species *i* in competition with the rest of the species *j*, and determines the shape of the functional responses. *D* is set to 1. [40].

### B. Synthetic invasion

To explore the network-level containment of single-species synthetic deployment, we rely on well established concepts from ecological invasions [41, 42]. The core idea is to take advantage of a resident species (likely to be a microorganism), and introduce a small genetic modification that confers some kind of advantage. Here for simplicity we assume that the introduced (synthetic) strain has a higher replication rate than the original wild-type (WT). The question we want to answer is what is the impact of these introduced species on the species composition and population abundances, and demonstrate that there exist controlled framework which engineers can rely on in order to assure safer synthetic interventions on ecosystems.

The bioengineering scheme thus involves starting from a stable community formed by *S* species. This community is obtained after simulating the dynamics from equation (1) starting from a small initial population *X_i_* (0) = *k_i_*/100 for all species. After a transient time *τ*_1_, the community trajectory is followed until it settles into a new equilibrium defined by per-iteration community change reduced below a threshold value (less than 10^−4^ movement in Euclidean space from the previous state). At that point we keep all the species such that *x_i_*(*τ*) > *θ* = 10^−5^ (otherwise, are removed) [36]. We thus have a stable community with the matrix

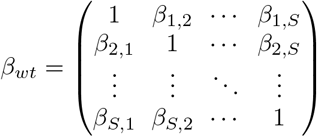

That is replaced by the “synthetic” community matrix, namely

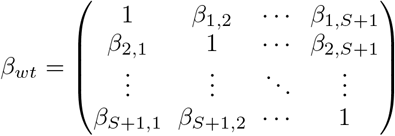

with *ρ*_*S*+1_ > *ρ_S_*. With an initial concentration of 0.1.

The goal now is to analyse the impact of this synthetic manipulation on the new community, and to weight the impact on the resident community. In particular, the success or failure of the invader and the number of extinct residents are determined and subsequently compared to the previous host community. To achieve a robust statistical analysis, core communities of different pool sizes are randomly generated. Invasions to each initial pool community are simulated over 100 runs (all of them with randomized community matrix and randomized replication rates). Constituting 100 experiments with different community interactions, same number of community members and different invaders. Thus we get 100 different outcomes for each species pool size. Additionally, the same numerical procedure is repeated 20 replicas and finally averaged.

## III. RESULTS

### A. Mean field theory

Consider first the synthetic invasion with a linear functional response in equation (1) (being *x_w_* the chosen wild type and *x_s_* the synthetic one). As defined above, we have now *β_w_j* = *β_sj_* (with *w* ≡ *S* + 1) but having an increased replication rate, i.e *ρ_s_* = *ρ_w_* + 0.1*ρ_w_*. Given that the new species is very close to its wild type ancestor, we can expect them to compete and sometimes exclude each other, perhaps generating extinction cascades. Can community-level properties act against such negative effects?

Before we explicitly explore the full multispecies problem, let us first consider a mathematical approach based on a mean-field approximation, where the exact nature of network correlations is ignored. Despite the high-dimensional nature of the problem, which usually makes analytic results difficult to obtain, the particular nature of the bioengineering design proposed here allows to make significant predictions.

Let us first consider the equilibrium state for the wild type in the non-invaded, resident (*R*) initial community, I. e.

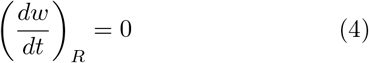

based on equation (1), this reads:

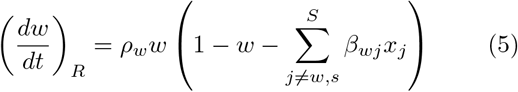

And thus the original equilibrium state was

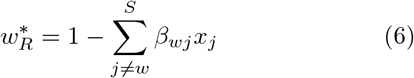

Since the synthetic strain is obtained from a wild type resident, there are some assumptions that can be made. As discussed above, the synthetic organism will share the community matrix values of a predefined wild type resident. In this case, we can write down -for our linear case-the following equations for the wild type (*w*) and synthetic (*s*) populations. Specifically we can see how the new equilibria as obtained from

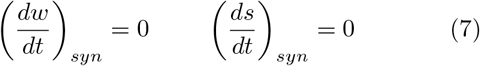

Can be approximated for the synthetic (*syn*) community. It is easy to see that the new equations for these two populations will read

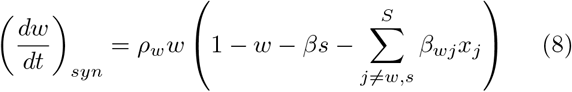

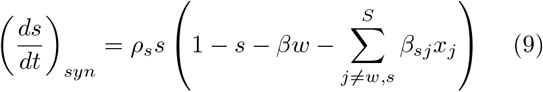

All equations share the same community interaction term

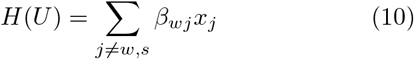

where *U* indicates the species set excluding both *w* and *s*. Now the new equilibrium state 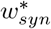 for the wild type population within the synthetic community can be easily related to the previous, non-manipulated valeues. Specifically, and since *β* = 1, we obtain:

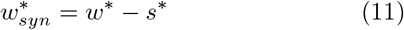

Which is our first prediction: the introduction of a faster-replicating strain derived from the original, resident one, will cause a drop in the later in an amount of the order of the introduced population. This is confirmed by numerical simulations of the full population dynamics, as shown in figure 2a. Here, once stabilization of the resident community has been achieved, the synthetic strain is introduced and two observations can be made. The first is that, as predicted from theory, both *w* and *s* persist. Secondly, the rest of the community barely reacts to the intervention.

**FIG. 2:**
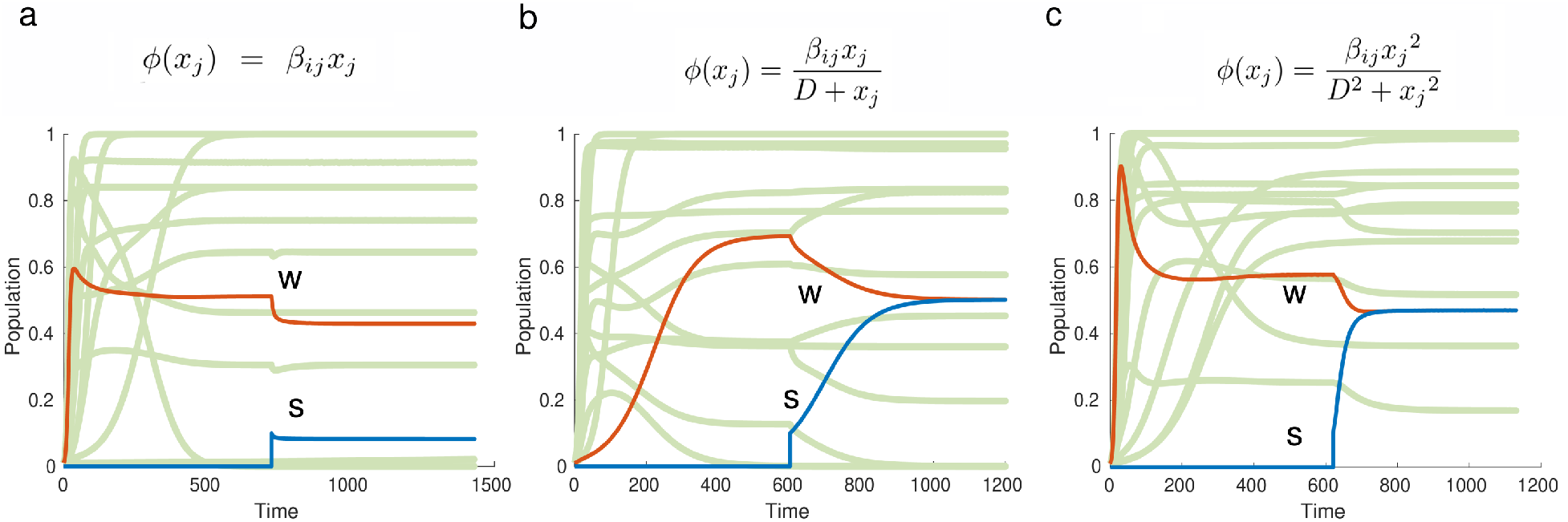
Population dynamics of single-species Terraformation. Three typical time series of the response to synthetic bioengineering are shown for the three functional responses considered here. Specifically, from left to right, we show the dynamics for linear, Holling I and Holling II, respectively. In all cases, we start from an initial ecosystem of 15 species at connectivity *C* = 0.08 and random *β* matrices. In orange we have the wild type and synthetic is represented in blue, whereas all other species are shown with the same color. The synthetic population is added when the initial community has reached its equilibrium state. Notice that, while for the linear scenario there is a drop in the wild type (similar to the steady population of the synthetic strain), for both Holling-I and Holling-II responses there is a convergence of both *w* and *s* to the same final state.

The same mathematical approach can be used to analyze the introduction of the synthetic strain for Holling non-linear responses, see equation. In this case, using again *β* = 1 and

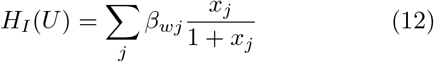

the new pairwise picture for Holling-I will be:

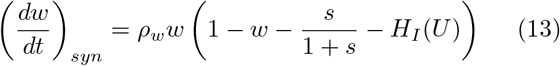

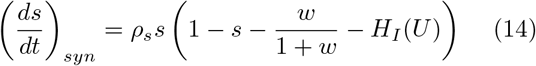

Now again if *w*\* = 1 – *H_I_*(*U*), and the new populations at equilibrium for the synthetic community are:

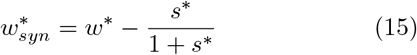

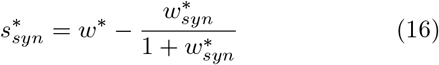

Finally, by substituting *w*\*, the following equality can be obtain an unexpected result, namely:

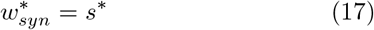

In other words, the resident strain used to engineer the synthetic one converge to exactly the same population value,

These results also hold for Holling-II responses, and the simulation results are displayed in figure 2(b-c). Here we can see that there is some re-arrangement of the population abundances in *x_k_* ∈ *U*, although all resident species are preserved, with rare exceptions (see next section). Because of the special nature of the synthetic ‘‘invader” the theoretical model indicates that the new community should in fact stabilize the introduced strain with little changes in the overall organization. In order to systematically test this suggestion, we need to perform a statistical analysis of ecosystem responses considering an ensemble of engineered communities.

### B. Synthetic community assembly

The previous results provide a basic prediction suggesting that the synthetic invader will be integrated by the resident ecosystem thus leading to species augmentation. However, system-level effects that are coarse-grained by the mean field approximations might modify our basic predictions. This is particularly relevant in relation with species richness: will less rich communities also behave in similar ways? In order to perform a statistical analysis of the likelihood of success of terraformation and its impact on community organization, we follow the classic work by Ted Case on alien invasions [41]. Specifically, using different levels of species diversity (*S*) wear at determining the predicted impact of the single-species terraformation. The statistics is obtained by performing the same class of perturbation as described above (fig. 2) where, for each *S* value, many terraformation simulations have been made using random sets of *β_ij_* and *ρ* parameters.

In general terms, an invasion event (including our case) can result in one of the next three potential outcomes. (i) *Invasion fails*: The invader is repelled, meaning that is unsuccessful at establishing within the host community. (ii) *Replacement*. One or more resident species of the host community get extinct when the invader succeeds at establishing. Finally, (iii) *Community Augmentation*: Invasion is successful while preserving all the previous resident species. In this case, the community absorbs the invader, growing in size by one [41].

Here we have considered both the standard random invader [41] and our synthetic scenario. For each *S*, 10^2^ different simulated ecosystems have been randomly generated and subsequently invaded to determine the frequency of each of the three possible outcomes described above. The whole process has been replicated for 20 runs to average the results. The results of our analysis are summarized in figure 3a-f where each outcome is represented as an ordered sequence (i. e. failure (white), replacement (black) and, augmentation (grey)) according to their frequency. In this way we can easily appreciate the effects of species diversity on the global outcome of the invasion event. Moreover, while Case’s study only involved linear interaction terms, we expand this by including the two other (non-linear) functional responses.

**FIG. 3:**
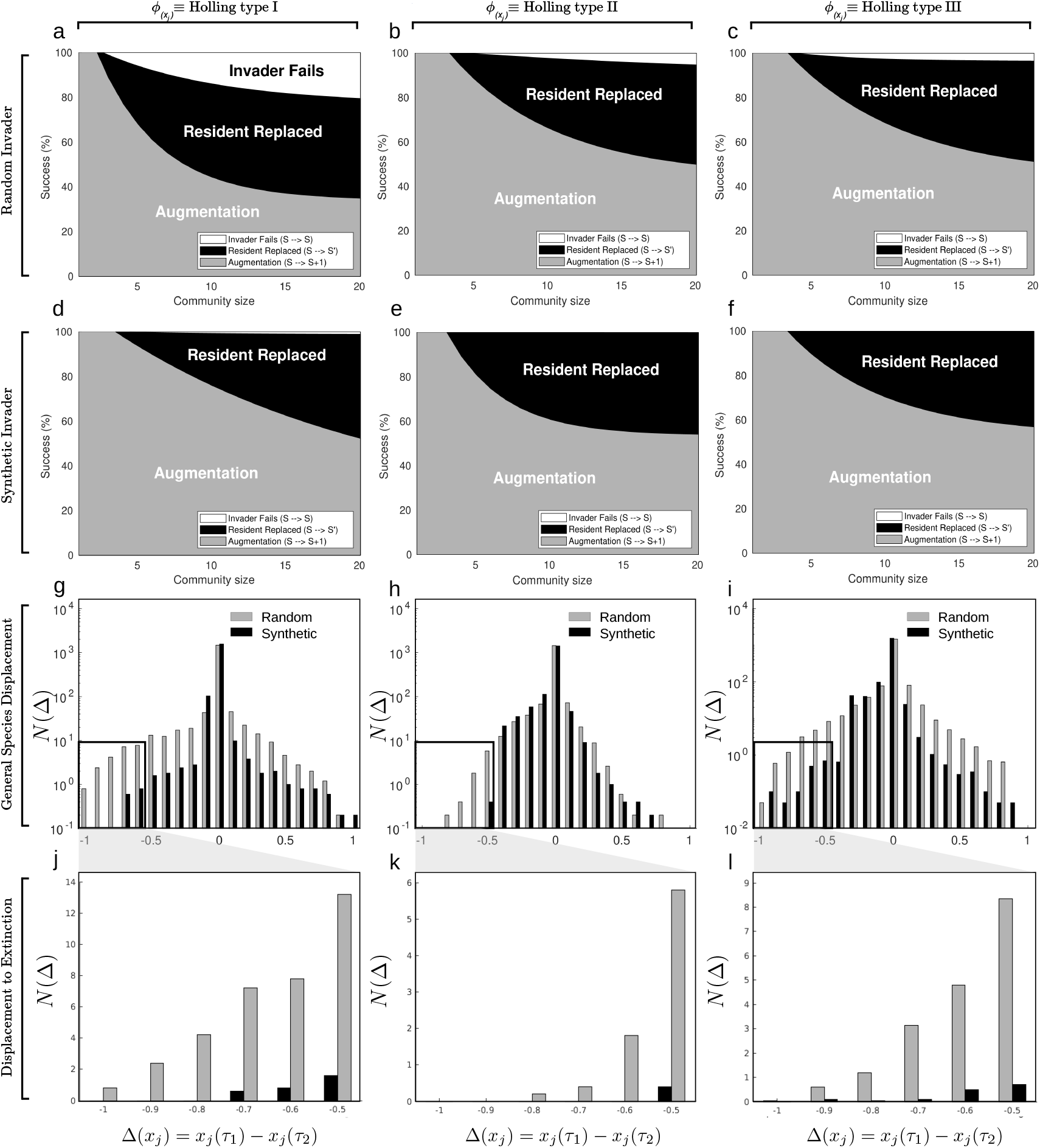
S-dimensional model results **(a-c)**, Failure, augmentation and replacement frequency of a classical random invader, as a function of community size and connectivity *C* = 0.08 as in [41]. Succes is the sum of augmentation and replacement. The same stands for synthetic invader in **(d-f)**. For each *S*, 10^2^ different simulated ecosystems randomly generated and invaded, replicated for 20 runs to average the results. In **(g-i)**, loss and gain of the species after the invasion process. Represented as displacement from −1 to 1, being −1 species that suffer maximum decay and 1 maximum gain. 0 displacement, means [−0.05 0.05]. For *S* = 15, 10^2^ ecosystem invasions, replicated 20 runs to average the results. Each time a given species from the community experience a change in their population abundance after invasion, a case accumulates in the graph. **(j-l)**, zoom on the left tail of the **(g-i)** distributions, using linear scales. All *β_ij_* are generated from a uniform distribution *U*[0,1].

For the linear interaction, while Case’s model revealed three possible phases for the (fig. 3a), including the failure of the invader to establish, the synthetic invader always succeeds, thus showing only two possible outcomes: augmentation or replacement of a resident species. As diversity grows, the likelihood of replacement increases for the linear case, while it saturates for the nonlinear functional responses (fig 3, b-c and e-f) while the probability that the invader fails shrinks for the random invader (b-c) and vanishes for the synthetic one (e-f).

While the previous plots provide a coarse view of the diversity-related responses to community invasion. Due to the blurred nature of the replacement category, further statistical measures are needed to determine the actual impact of introduced species. One way of doing so is to estimate the population displacement for each population,

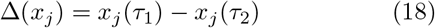

associated to the reconfiguration event post invasion, accumulated through all the statistical trials for a *S* = 15 community. Here *τ*_1_ is the last time step before the introduction of the synthetic, once the resident population has reached its stable state, and the final population is computer after approx *τ*_2_ steps (see methods). In figures 3g-i a histogram of the number of species experiencing a displacement of size Δ ∈ [−1, +1]. In these plots, a logarithmic scale is used for *N*(Δ). It is immediately apparent that most events had none or little displacement, with marked differences in the negative side of the Δ axis, associated to population loss.

While random invaders (grey bars) can cause considerable displacements and species extinctions (as Δ < −1 events are possible) in the linear interaction model, this is not the case for the synthetic displacement profile (black bars) which are much less common (in some values in an order of magnitude less) and seldom cause extinctions. A similar pattern is obtained for Holling-I response (fig. 3h,k) and less marked for Holling II.

A close inspection of the left tail of the distributions further improves our picture of the distinct nature of the invasion processes. As shown in fig 3j-l, now using linear scales, single-species bioengineering has a much reduced negative impact on ecosystem networks. Here we zoom into the interval [−1, −1/2] of population decays that could easily trigger extinctions.

## IV. DISCUSSION

The question of the potential, unintended consequences of deploying bioengineered organisms in the wild has been a recurrent topic. Due in part to the comparison with harmful invasion events, it has been too often assumed (with no further scientific evidence) that modified strains will necessarily involve negative, perhaps catastrophic consequences. Is that the case? The honest answer to that question must necessarily acknowledge our ignorance. We do not know. But knowing the answer is becoming more and more relevant for a number of reasons. One is that the rapid development of synthetic biology necessarily asks for understanding the potential consequences of leaked microorganisms. Secondly, this kind of engineering is being developed outside the academic boundaries, with many entrepreneurs and amateurs building genetic circuits in their domestic labs. Finally, and most important, the rapid pace of ecosystem deterioration due to global warming asks for novel ways of approaching the problem of how to protect fragile ecosystems from degradation and collapse [19, 30].

As it occurs with most relevant questions in ecology, simple models can help finding tentative answers or, to the least, a rationale for the expected patterns and processes that we want to understand. With this aim, we have presented here a very simple mathematical model of ecosystem engineering that includes the minimal class of intervention that can be designed. As suggested in previous work on ecosystem terraformation [19, 29] one potentially successful way of modifying ecosystems could involve the modification of an actual member of the resident community that would then be introduced as a “synthetic” strain incorporating some minimal genetic modification. Here we have limited ourselves to a simple change that allows the modified individuals to replicate faster than the wild type strain from which they have been designed. The rest of the interactions remain the same. Under these idealized circumstances, one could conjecture that the introduced, faster-replicating synthetic species would overcome (and displace) the original strain and perhaps alter the community in unintended ways. This is not the case. Instead, our model actually reveals that synthetic strains should be expected to establish within the resident community, coexisting with the wild type. Actually, when nonlinear responses are taken into account, we predict a closely similar population levels for both strains.

How robust are our model predictions? By considering the classical analysis on community assembly [41] we have been able to compare those results with our synthetic invasion mechanism. Future work should consider several extensions and more general assumptions, such as relaxing the symmetry constraints on the interaction matrix or the introduction of mutualistic effects, which are known to be relevant in microbial soil communities [43, 44]). Similarly, resource-consumer models would be also useful to test the role played by the engineered strain on available resources. Additionally, one can move beyond the single-species picture and consider instead the multi-species dissemination of mobile DNA (see [45] and references cited). Finally, these in silico experiments could be implemented in microbial microcosms [46, 47] as an intermediate step towards the application of terraformation strategies to real case studies [29].

## V. ACKNOWLEDGMENTS

The authors thank the Complex Systems Lab members for fruitful discussions. Special thanks to Blai Vi-diella, Josep Sardanyes, Fernando Maestre and Miguel Berdugo. V.M. thanks Jordi Piñero, Guim Aguadé, Luis Seoane and Thomas Wake for the inspiring conversations at the Santa Fe Institute. This work was supported by the PR01018-EC-H2020-FET-Open MADONNA projectBotén Foundation by Banco Santander through its Santander Universities Global Division, the Spanish Ministry of Economy and Competitiveness, grant FIS2016-77447-R MINECO/AEI/FEDER, an AGAUR FI-SDUR 2020 grant, and the Santa Fe Institute, where most of this work was done.

